# Mitochondrial DNA variation across 56,434 individuals in gnomAD

**DOI:** 10.1101/2021.07.23.453510

**Authors:** Kristen M. Laricchia, Nicole J. Lake, Nicholas A. Watts, Megan Shand, Andrea Haessly, Laura Gauthier, David Benjamin, Eric Banks, Jose Soto, Kiran Garimella, James Emery, Genome Aggregation Database Consortium, Heidi L. Rehm, Daniel G. MacArthur, Grace Tiao, Monkol Lek, Vamsi K. Mootha, Sarah E. Calvo

## Abstract

Databases of allele frequency are extremely helpful for evaluating clinical variants of unknown significance; however, until now, genetic databases such as the Genome Aggregation Database (gnomAD) have ignored the mitochondrial genome (mtDNA). Here we present a pipeline to call mtDNA variants that addresses three technical challenges: (i) detecting homoplasmic and heteroplasmic variants, present respectively in all or a fraction of mtDNA molecules, (ii) circular mtDNA genome, and (iii) misalignment of nuclear sequences of mitochondrial origin (NUMTs). We observed that mtDNA copy number per cell varied across gnomAD cohorts and influenced the fraction of NUMT-derived false-positive variant calls, which can account for the majority of putative heteroplasmies. To avoid false positives, we excluded samples prone to NUMT misalignment (few mtDNA copies per cell), cell line artifacts (many mtDNA copies per cell), or with contamination and we reported variants with heteroplasmy greater than 10%. We applied this pipeline to 56,434 whole genome sequences in the gnomAD v3.1 database that includes individuals of European (58%), African (25%), Latino (10%), and Asian (5%) ancestry. Our gnomAD v3.1 release contains population frequencies for 10,850 unique mtDNA variants at more than half of all mtDNA bases. Importantly, we report frequencies within each nuclear ancestral population and mitochondrial haplogroup. Homoplasmic variants account for most variant calls (98%) and unique variants (85%). We observed that 1/250 individuals carry a pathogenic mtDNA variant with heteroplasmy above 10%. These mitochondrial population allele frequencies are publicly available at gnomad.broadinstitute.org and will aid in diagnostic interpretation and research studies.

## INTRODUCTION

The genetic material of human cells is contained in the nucleus and mitochondria. The mitochondrial genome (mtDNA) is a circular molecule of 16,569bp containing 37 genes that encode 13 proteins, 22 tRNAs, and 2 rRNAs (Anderson et al. 1981), all essential to mitochondrial electron transport and energy homeostasis. Depending on the tissue, human cells contain hundreds to thousands of copies of mtDNA. Because the maternally-inherited mtDNA does not recombine and exhibits a ten times greater rate of polymorphism than nuclear DNA, it has been extremely useful in tracking human biogeography (Brown et al. 1979; Cann et al. 1987; Cavalli-Sforza 1998).

Pathogenic variants in the mtDNA (Lott et al. 2013; Gorman et al. 2016) account for ~80% of adult-onset and ~20% of pediatric-onset mitochondrial disease (Gorman et al. 2015, 2016). Pathogenic mtDNA variants can cause disease at homoplasmy or when heteroplasmy rises to high levels (Craven et al. 2017). These latter disorders are particularly challenging to diagnose since pathogenic variants can sometimes be observed at lower heteroplasmy levels and even absent in blood versus affected tissue and can decrease over time (Grady et al. 2018). For both homoplasmic and heteroplasmic variants, distinguishing those that are pathogenic from those that are benign is a challenge.

Population frequency data are extremely helpful for the clinical interpretation of variants of uncertain significance (VUS) (McCormick et al. 2020). Until now, mtDNA variants have not been included in most large population databases of genomic variation such as the Exome Aggregation Consortium (ExAC) (Lek et al. 2016), the Genome Aggregation Database (gnomAD) (Karczewski et al. 2020), and the BRAVO server (https://bravo.sph.umich.edu). Instead, four specialized databases provide mtDNA population frequencies across humans: (i) MITOMAP (Lott et al. 2013) provides population frequencies from GenBank (which has heterogeneous data quality and is known to include individuals with disease); (ii) HmtDB provides population frequencies from GenBank and from mitochondrial disease patients (Clima et al. 2017); (iii) MSeqDR compiles population frequency data including MITOMAP, HmtDB, and GeneDx (Shen et al. 2018); (iv) HelixMTdb uses proprietary exome technology on saliva samples to report homoplasmic and heteroplasmic variants for nearly 200,000 individuals (mainly of European ancestry), despite relatively low (~180x) mean mtDNA coverage that can make it difficult to call heteroplasmic variants (Bolze et al. 2020). The first three mtDNA databases report only homoplasmic variants.

The Genome Aggregation Database (gnomAD) is a widely-used resource of human genetic data developed by an international consortium who have aggregated whole genome sequence (WGS) data from large-scale sequencing projects including The Cancer Genome Atlas (TCGA), the Centers for Common Disease Genomics (CCDG), the Genotype-Tissue Expression project (GTEx), the Trans-Omics for Precision Medicine (TOPMed), and the National Heart, Lung, and Blood Institute (NHLBI). The data in gnomAD are analyzed jointly using the same pipeline, and are depleted for Mendelian or severe pediatric diseases as well as for cryptically related individuals, allowing for the computation of accurate and high-quality population allele frequencies. Summary gene and variant metrics are made public for a range of diverse ancestral groups, including individuals of African and African-American, Amish, Latino and admixed American, Ashkenazi Jewish, East Asian, Finnish, non-Finnish European, Middle Eastern, and South Asian descent. gnomAD v2 contains 125,748 exomes and 15,708 genomes aligned to the GRCh37 human reference. gnomAD v3 contains 76,156 WGS samples aligned to the GRCh38 human reference and includes cohorts derived from controls and biobanks (~16.5K), TOPMed (~35.7K), non-pediatric neurological disease cohorts (~8.7K, including individuals with schizophrenia, Alzheimer’s disease, migraines, bipolar, and affective and psychotic disorders), and others. The resource has been widely used for both basic and clinical research, with ubiquitous adoption in clinical genetic diagnostic pipelines worldwide, but analysis of the mtDNA has not been included until now.

The main challenge for mtDNA variant calling from WGS data is to distinguish low heteroplasmy variants from sample contamination, sequencing errors, and misalignment. Specifically, misalignment from nuclear sequences of mitochondrial origin (NUMTs) is particularly problematic because the reference genome contains several hundred ancient NUMTs (Li et al. 2012) and hundreds of “polymorphic NUMTs” not present in the reference genome (Dayama et al. 2014), including rare instances of large, tandemly repeated mega-NUMTs (Lutz-Bonengel et al. 2021). In addition, the circular mtDNA molecule can present alignment challenges, and many alignment algorithms show a drop of coverage at the artificial ends of the linearized chrM chromosome. Since nuclear variant pipelines are not suitable for mtDNA variant calling, the mtDNA has not been routinely analyzed by many WGS projects.

Multiple tools exist to call mtDNA variants. Tools such as mtDNA-Server (Weissensteiner et al. 2016a), MToolBox (Calabrese et al. 2014), and mity (Puttick et al. 2019) have been designed specifically to call heteroplasmic and homoplasmic variants. mtDNA-Server specifically identifies contamination, and MToolBox aims to avoid misalignment of NUMTs in the reference assembly but cannot avoid polymorphic NUMTs. Other tools not specifically designed for mtDNA can be adapted to call heteroplasmic variants, such as GATK Mutect2 (Benjamin et al. 2019), which was originally designed to identify sub-clonal variants in cancer. Many of these tools are easy to run; however, by themselves, they do not address issues such as contamination and false-positives from misalignment.

Here, we aimed to accurately call mitochondrial variants in gnomAD WGS samples. To achieve highly accurate calls with few false positives and false negatives, we excluded samples we found to be particularly prone to NUMT misalignment (having few mtDNA copies per cell), samples likely to have cell line artifacts, and samples showing contamination from other samples. Next, we report only variants with heteroplasmy ≥ 10% because we observed a significant fraction of NUMT-derived false positives below this threshold. Thus gnomAD v3.1 conservatively reports variants above 10% heteroplasmy in 56,434 samples.

## RESULTS

### mtDNA coverage varies across cohorts in gnomAD

WGS provides even coverage across the mtDNA for all 70,375 gnomAD v3 samples available for analysis (Fig. 1A). However, we find that mtDNA coverage, as well as mtDNA copy number per cell (mtCN), vary widely across gnomAD cohorts, independent of nuclear coverage (Fig. 1B–E). This variation likely depends on source material (e.g. blood, buffy coat, cell line, tissue) and DNA extraction protocol; however, such annotations are available only for a subset of samples. A typical blood sample with 30x nuclear coverage shows ~2700x mtDNA coverage. The high mtDNA coverage in WGS enables detection of candidate heteroplasmic and homoplasmic variants. We estimate mtCN as 2m/n where m is mean mtDNA coverage and n is median nuclear coverage. As expected, mtCN varies by source material. Blood samples show two distinct peaks (median 40 for TOPMED COPD and 207 for NHLBI cohorts) possibly associated with DNA extraction kits or blood cell types collected (Fig. 1E). Cell lines typically have 500-1200 mtDNA copies per cell. A small number of samples with outlier mtCN > 2000 are derived from tissue samples such as heart, adrenal, and kidney.

**Figure 1.**
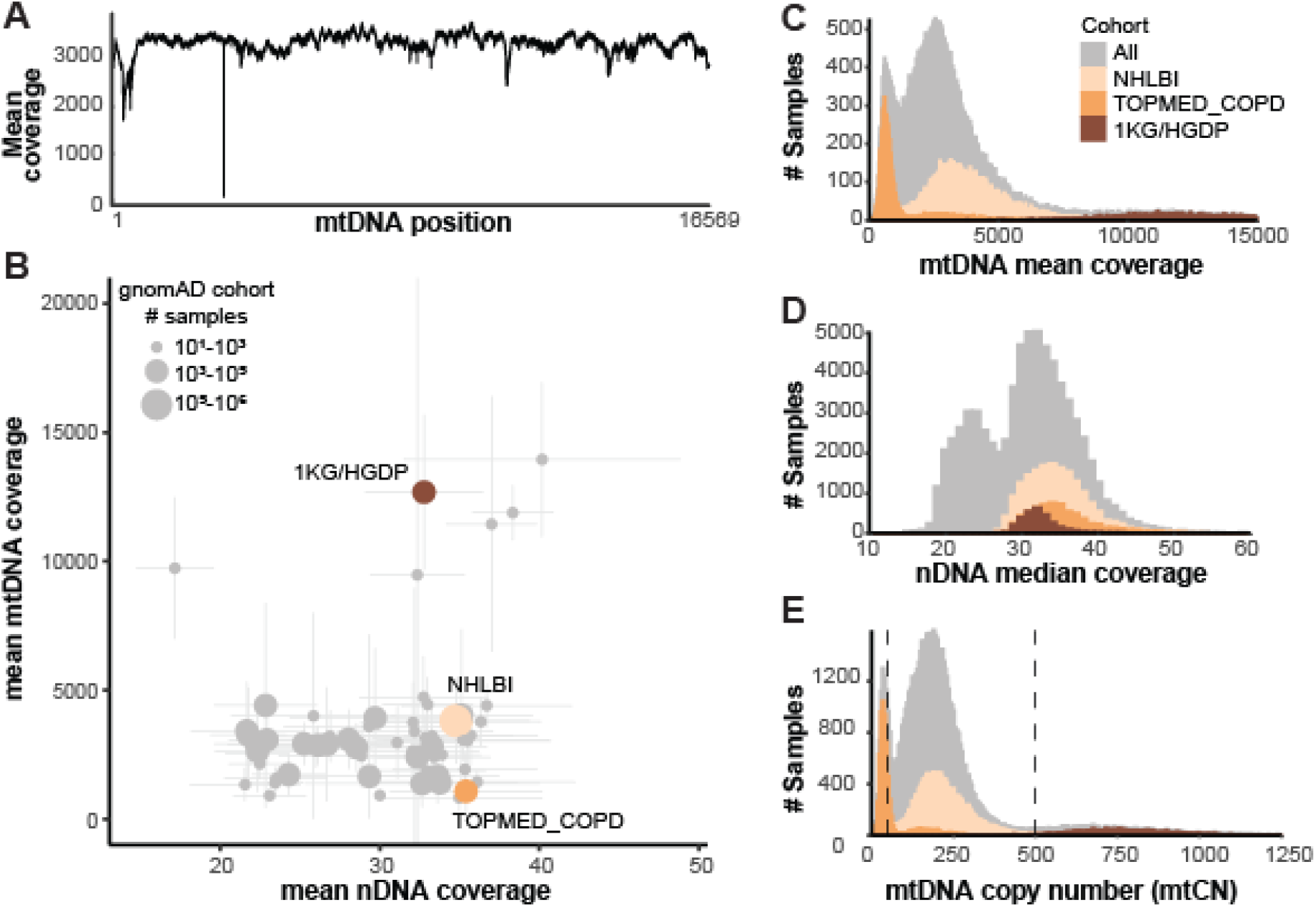
Coverage statistics for 70,375 gnomAD WGS samples. **(A)** Per-base mean depth of coverage across mtDNA, with coverage dips at positions 303-315 and 3107 due to homopolymeric tract and chrM reference deletion respectively. **(B)** For each cohort within gnomAD, scatter plot shows the mean nuclear (nDNA) and mtDNA coverage +/− standard deviation. Three example cohorts are shown in color: 1000 Genomes and Human Genome Diversity Project cell lines (1KG/HGDP), NHLBI, and TOPMed Chronic Obstructive Pulmonary Disease (TOPMED COPD). **(C)** Histogram shows mean mtDNA coverage for all samples, and overlaid histograms show three selected cohorts (806 outliers with coverage 15000-97000 excluded). We note mean and median (not shown) mtDNA coverage statistics are extremely similar. **(D)** Histogram shows median nDNA coverage for all samples, and overlaid histograms show three selected cohorts (84 outliers with coverage 60-94 excluded). **(E)** Histogram shows mtDNA copy number per cell (2*mean mtDNA coverage/ median nDNA coverage) for all samples, and overlaid histograms show three selected cohorts (223 outliers with mtCN 1250-7000 excluded). Only samples with mtCN 50-500 (dashed lines) were included in the released mtDNA call set (56,434/70,375).

### Pipeline for mtDNA variant calling in individual samples

We developed a high-throughput GATK pipeline to call homoplasmic and heteroplasmic variants in mtDNA from whole genome sequence data (Fig. 2A). WGS was aligned to the reference genome using BWAmem (Li 2013). Only mate-pairs with both reads mapping to chromosome M were used for variant calling, after excluding duplicate pairs. Variants were called using the GATK Mutect2 variant caller (Benjamin et al. 2019), parameterized via a specific “mitochondria mode” designed to account for high coverage and potential low-heteroplasmy variants. To call variants in the control region that spans the artificial break in the circular genome (coordinates chrM:16024-16569 and chrM:1-576), we extracted all chrM reads and realigned them to a mtDNA reference that was shifted by 8,000 bases, called variants on this shifted alignment, and then converted coordinates back to their original positions. Variants showing weak evidence or strand bias were then filtered. Variant allele fraction (VAF) was calculated as the fraction of alternate reads to total reads for each variant and sample. We denote variants with VAF 0.95-1.00 as homoplasmic or near homoplasmic, and variants with VAF < 0.95 as heteroplasmic.

**Figure 2.**
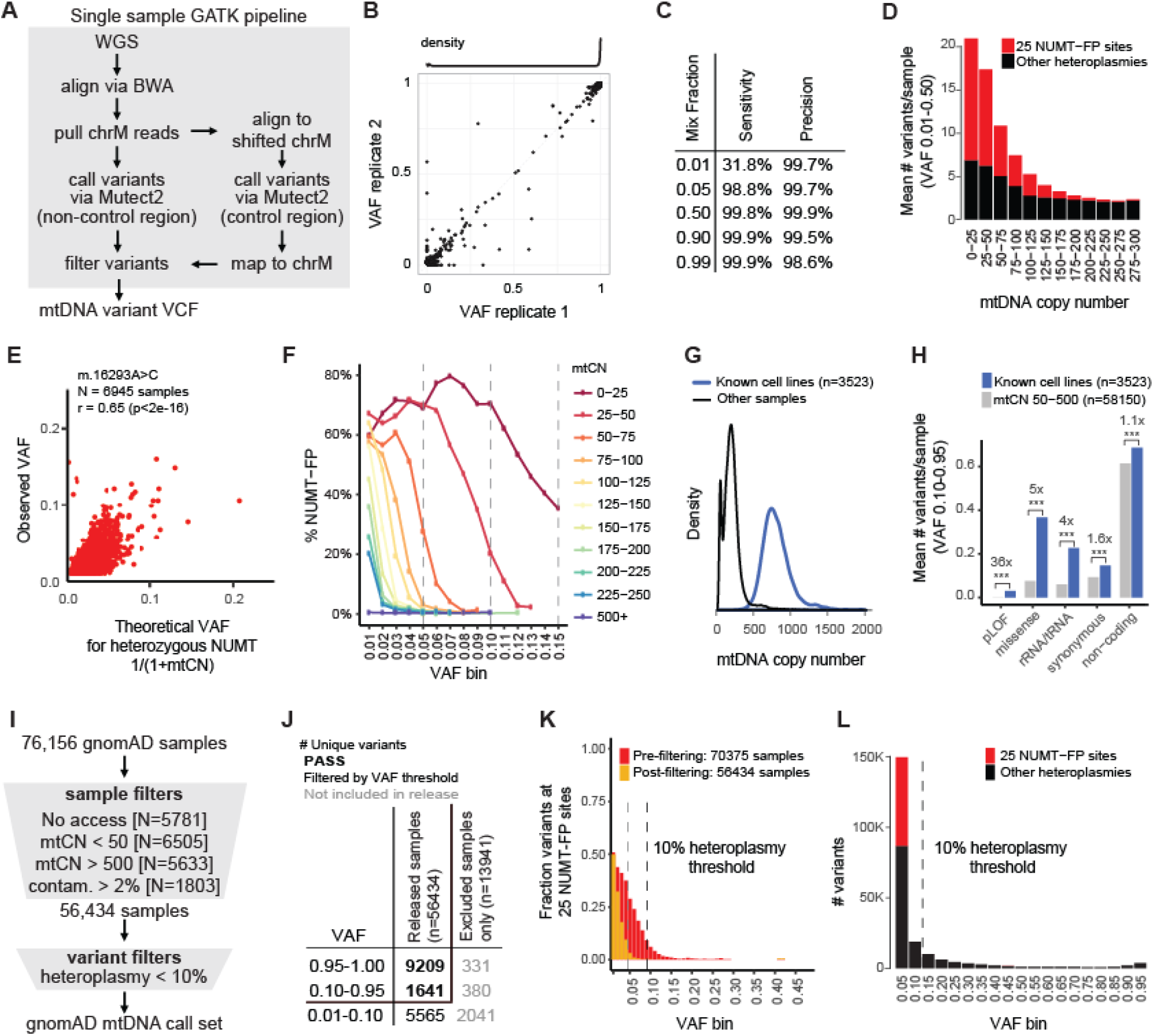
mtDNA call set is designed to exclude NUMT-derived false positives (NUMT-FPs), cell line artifacts, and contaminants. **(A)** Schematic shows GATK pipeline for calling mtDNA variants in single WGS samples. The control region spans the artificial break in chromosome M sequence. **(B)** Reproducibility of GATK pipeline on 91 WGS replicate samples shows 99.3% concordance of calls (2533/2551), and density plot at top shows 87% variants are homoplasmic. **(C)** Accuracy of single-sample pipeline in samples with mtCN > 500 based on “in silico” mixing data. **(D)** Barchart shows that the mean number of putative heteroplasmies per sample depends on mtDNA copy number (mtCN), as does the subset occurring at 25 validated NUMT-FP sites (red). **(E)** Scatterplot shows the observed VAF for a single NUMT-FP (m.16293A>C) across 6,945 samples versus the theoretical VAF if the NUMT were heterozygous and all reads misaligned to chrM. (**F**) Plot shows VAF levels for NUMT-FP sites decrease with mtCN (colored lines). Y-axis indicates the percent of detected variants that occur at 25 NUMT-FP sites. **(G)** Density plot shows mtCN for known cell lines and all other samples. **(H)** Barplot shows that known cell lines have increased number of heteroplasmic variants in all categories compared to samples with mtCN 50-500 (enrichment shown with *** indicating p-value < 1e-5 based on Fisher’s Exact test); pLOF indicates predicted loss-of-function. **(I)** Schematic shows steps for combining and filtering single-sample variant calls into the gnomAD mtDNA call set, designed to exclude NUMT-derived false positives, cell line artifacts, and contaminants. (**J**) Number of unique variants that pass filters (bold black) versus those filtered out based on VAF (black) or not released (gray). The 19,167 variants are partitioned into mutually exclusive categories, e.g. VAF 0.10-0.95 excludes variants also detected VAF 0.95-1.00. **(K)** For each VAF level, barchart shows the fraction of variants at 25 NUMT-FP sites before sample filtering (red) or after filtering (orange, shown overlaid). **(L)** Histogram of VAF (after sample filtering) shows that below 10% VAF, there are a large number of variants and a substantial fraction present at 25 validated NUMT-FP sites (red). X-axis label indicates upper bound of VAF bin.

We assessed the reproducibility of our pipeline using 91 samples for which replicate WGS was available (Fig. 2B). We observed 99.3% concordance for all variants with VAF ≥ 0.01, where concordance is defined as the number of variants detected in both samples / number of variants detected in either sample. Some of the highly discordant calls were derived from cell lines, which may have accumulated mutations over the times sampled.

To assess sensitivity and precision at different heteroplasmy levels, we created in silico mixtures of samples (Fig. 2C) to model variants at specific VAF levels (0.01, 0.05, 0.50, 0.90, 0.99). We mixed truth data from cell line NA12878 with each of 22 African-haplogroup samples to increase the total number of variants (1,200 variants at 286 positions, including 8 indels). For VAF ≥ 0.05, we observed excellent sensitivity (99-100%) and precision (98.9-99.7%), where sensitivity indicates the percent of true variants that are detected, and precision indicates the percent of detected variants that are true positives. Sensitivity dropped to 32% for variants at 0.01 VAF. Compared to the mtDNA-Server algorithm, GATK Mutect2 had higher precision at all heteroplasmies, similar sensitivity for VAF 0.05-0.99, but reduced sensitivity for VAF 0.01 variants (Fig. S1).

We note that this in silico approach uses cell lines and does not account for possible NUMT-misalignment, which we show is very problematic for samples with low mtDNA copy number.

### NUMT-derived false positives anti-correlate with sample mtDNA copy number and VAF

When we applied this variant calling pipeline to 70,375 available whole genomes in gnomAD v3.1, we observed the number of candidate heteroplasmies per sample was highly dependent on the sample mtCN (Fig. 2D). This observation was consistent with false positives derived from NUMT-misalignment. Theoretically, a misaligned heterozygous NUMT will have VAF approximately 0.5n/(0.5n+m) or 1/(1+mtCN) where n is nuclear coverage and m is mtDNA coverage and mtCN=2m/n. We observed several dozen common variants whose VAF correlated with 1/(1+mtCN) (e.g. m.16293A>C, Fig. 2E), and were often linked in *cis* to each other (Table S1). We hypothesized that these were derived from polymorphic NUMTs, i.e., NUMTs present in some individuals but not in the reference genome assembly. We validated two polymorphic NUMTs using long-read PacBio data: numtA (871bp insertion from chrM:12361-13227 into chr21:9676568), and numtB (536bp insertion from chrM:16093-chrM:59 into chr11:49862017). When misaligned to the reference mitochondrial genome, these two NUMTs together yielded 25 common false positive calls that we term NUMT-FP (Fig. S2, Table S1). Some of the false positives were properly filtered out by strand bias, but others passed our variant filters. Using unfiltered variant calls, we estimate numtA and numtB are each present in ~40% of individuals in our data set (Table S1, Fig. S2).

Next, we aimed to estimate the extent of NUMT-misalignment and how it relates to sample mtCN and VAF. As a lower bound we can assess the percent of variants at each VAF level located at these 25 NUMT-FP sites (requiring each NUMT to be supported by at least two NUMT-FP per sample). As expected, the 25 NUMT-FPs were more problematic for samples with low mtCN, and for variants with low VAF (Fig. 2F). Samples with extremely low mtCN (< 50) showed substantial NUMT-FP exceeding 0.15 VAF. For mtCN 50-75, there were detectable NUMT-FP variants up to 0.10 VAF. For samples with mtCN 75-100, there were substantial NUMT-FP up to 0.05 VAF. For samples with mtCN > 500, there were almost no NUMT-FP with VAF ≥ 0.01. As expected, shorter WGS insert sizes also cause greater misalignment (Fig. S3). The true extent of NUMT-derived false positives is likely to be much higher, since this analysis considers only two common NUMTs, whereas there are hundreds of known polymorphic NUMTs (Dayama et al. 2014) and hundreds of NUMTs yet to be identified.

Given these large numbers of false positive calls for variants with VAF < 0.10, for the initial release we chose to exclude samples with mtCN < 50 and to report only variants with VAF ≥ 0.10 as we have greater confidence that such variants represent genuine heteroplasmies and not NUMT-derived false positives (Fig. 2F).

We note that misalignment due to NUMTs not only causes false positive calls at low VAF, but also can cause truly homoplasmic variants to appear heteroplasmic, with the reference alleles derived from the misalignment of a NUMT. Because of this, we term all variants with VAF 0.95-1.00 as “homoplasmic” or “near-homoplasmic.”

### Cell lines show excess deleterious heteroplasmies

While not all samples have annotations of source material, the 3,436 known cell lines account for the majority of the 5,633 samples with mtCN > 500 (Fig. 1E, 2G). Samples annotated to be cell lines show significantly elevated numbers of heteroplasmies (VAF 0.10-0.95), with a particular excess of potentially deleterious variants (loss of function, missense, tRNA, and rRNA) compared to synonymous and non-coding variants (Fig. 2H). These data show that cell lines accumulate mutations and suggest that deleterious mtDNA variants may be tolerated in cell culture.

### Filtering gnomAD samples and variants

We performed stringent filtering of samples to create a high-quality mtDNA variant call set (Fig. 2I). Specifically we excluded: (1) 6,505 samples with mtCN < 50 to avoid excessive misalignment due to NUMTs; (2) 5,633 samples with mtCN > 500 since these were primarily cell lines and enriched with what appear to be cell culture derived variants; (3) 1,803 samples with contamination exceeding 2% based on estimates from the nuclear DNA or mitochondrial DNA, since samples with low nuclear contamination can still have substantial mtDNA contamination. No data from these excluded samples are provided in the release.

In the remaining 56,434 samples, we conservatively report only mtDNA variants with VAF ≥ 0.10 (Table S2). 10,850 unique variants pass our thresholds whereas the remainder (including variants VAF 0.01-0.10, Fig. 2J) are available as filtered variants in download files and on the web portal but are plagued with false positives (Fig. 2K). Using just two validated NUMTs, we calculate a lower bound for NUMT-misalignment, which accounts for 42% of all variant calls VAF 0.01-0.05, 1% of all variants VAF 0.05-0.10, and virtually 0% of variants in other heteroplasmy bins (Fig. 2L). Variants enriched for false positives are annotated and highlighted in the web browser, e.g. using the “common_low_heteroplasmy” flag (variants detected at VAF 0.001-0.50 in > 56 individuals), “artifact_prone_site” filter, “indel_stack” filter, or “no pass genotype” filter (see Methods, Table S2). Since variants with VAF 0.10-0.95 have few false positives, we refer to these as heteroplasmies.

### mtDNA variants across 56,434 gnomAD samples

We release high confidence mtDNA variants for 56,434 samples that pass our quality control filters. These samples exhibited median 2,700x mtDNA coverage and 184 mtCN (Fig. S4). Overall, 8,793 of the 16,569 mtDNA nucleotides had a variant (53%) (Fig. 3A). We observed 10,850 unique variants, including 10,434 SNVs (96%) and 416 indels (4%), with SNVs being predominantly transitions rather than transversions (Fig. 3A). Of the 1.9M total variant calls, 98% were homoplasmic or near-homoplasmic and 2% were heteroplasmic (40,706 variant calls 10-95% heteroplasmy) (Fig. 3A, Fig. S4D). The 9,209 unique homoplasmic variants include known haplogroup markers (46%) as well a large number of rare variants. Homoplasmic variants showed a range of population frequencies (Fig. 3B–C).

**Figure 3:**
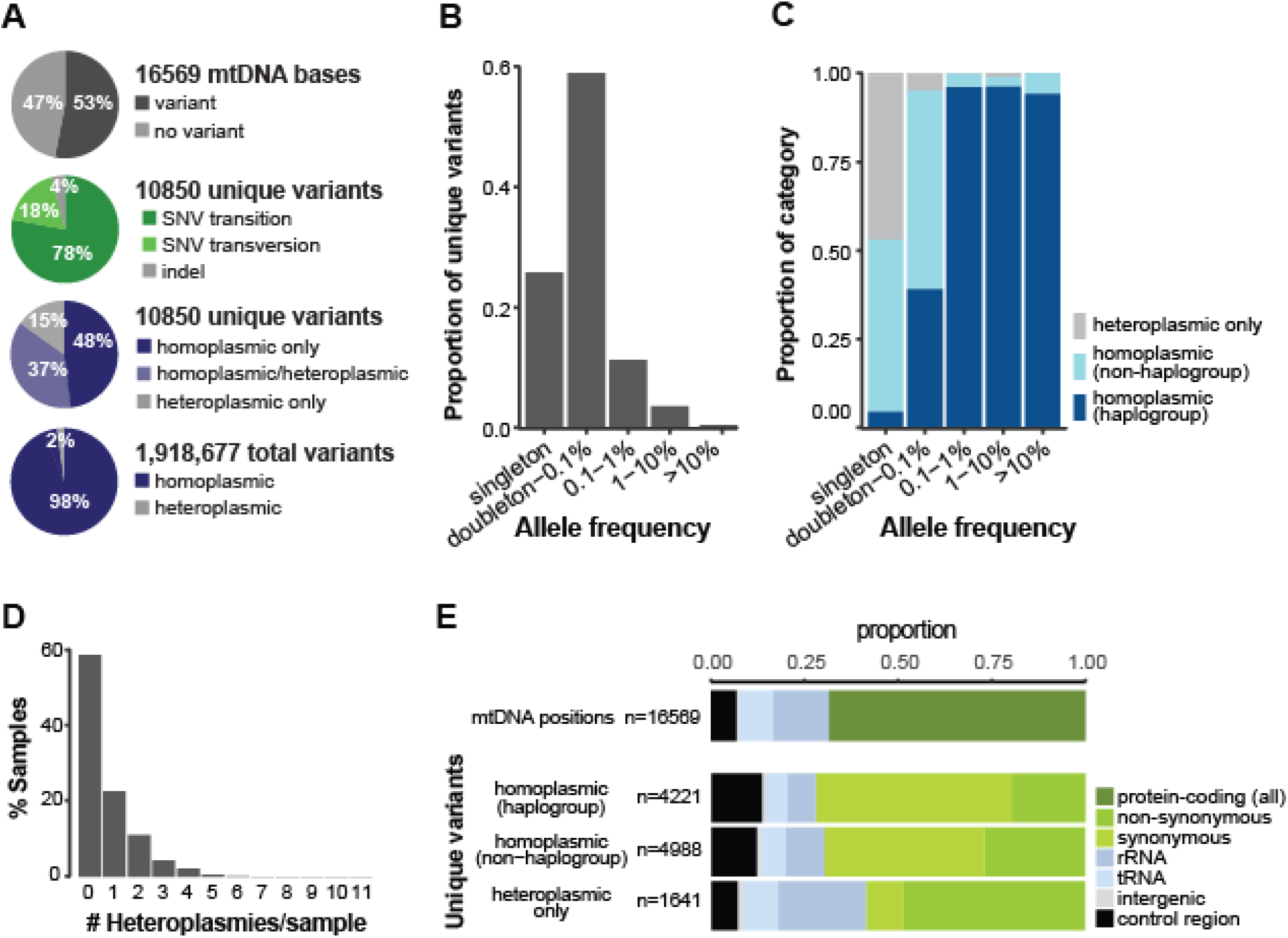
gnomAD mtDNA variant statistics. **(A)** Pie charts summarize statistics on mtDNA bases with variants, unique variants, and total variant calls. **(B)** Bar plot shows the proportion of unique mtDNA variants detected at different population allele frequencies in gnomAD v3.1. **(C)** Bar chart shows the proportion of variants that are observed only at 10-95% heteroplasmy (gray) or observed at homoplasmy (blue) including those that are known haplogroup markers in Phylotree (dark blue). **(D)** Histogram shows number of heteroplasmies per sample. **(E)** Stacked bar charts show the distribution of variant annotations in the entire mtDNA and for unique variants that are homoplasmic or only observed at heteroplasmy.

The majority of samples had no heteroplasmies (Fig. 3D). Moreover, the majority of heteroplasmies (32,386/40,706=80%) were variants that were detected at homoplasmy in other samples. Specifically, 5,205 (48%) unique variants were observed only at homoplasmy, 4,004 (37%) were observed both as homoplasmic and heteroplasmic, and 1,641 (15%) were observed only at heteroplasmy (Fig. 3A). Most unique variants observed only at heteroplasmy were found in only one or two samples (Fig. 3B–C). Variants observed only at heteroplasmy showed increased non-synonymous and RNA gene changes, whereas variants observed at homoplasmy showed higher prevalence of synonymous and non-coding variants (Fig. 3E).

### Haplogroups vs nuclear ancestry

Since mtDNA does not recombine and is inherited maternally, closely related mtDNA sequences have historically been grouped together in “haplogroups”. There are 5,184 haplogroups from diverse populations available in the Phylotree database (van Oven and Kayser 2009) and broadly associated with African, Asian, and European ancestry (Lott et al. 2013). Samples in gnomAD v3.1 spanned 61% of the haplogroups defined by Phylotree, and provide representation from 29/33 of the top-level haplogroups (missing L6, Q, O, S) (Fig. 4A). 46% of gnomAD v3.1 homoplasmic unique variants were known haplogroup markers, and 4250/4571 (93%) of all haplogroup markers were observed in the dataset, emphasizing the haplogroup and population diversity of the samples included in the current release. The mtDNA reference sequence in hg38, also known as the revised Cambridge Reference Sequence (rCRS), belongs to the European top-level haplogroup H (Andrews et al. 1999). Accordingly, the number of homoplasmic mtDNA variants per person in gnomAD increased as distance from the reference haplogroup in the phylogenetic tree increased, where individuals of European haplogroups typically had 0-50 variants, Asian haplogroups typically had 25-50 variants, and African haplogroups typically had 50-100 variants (Fig. 4B). By contrast, the number of heteroplasmic variants was similar across haplogroups (Fig. S5).

**Figure 4:**
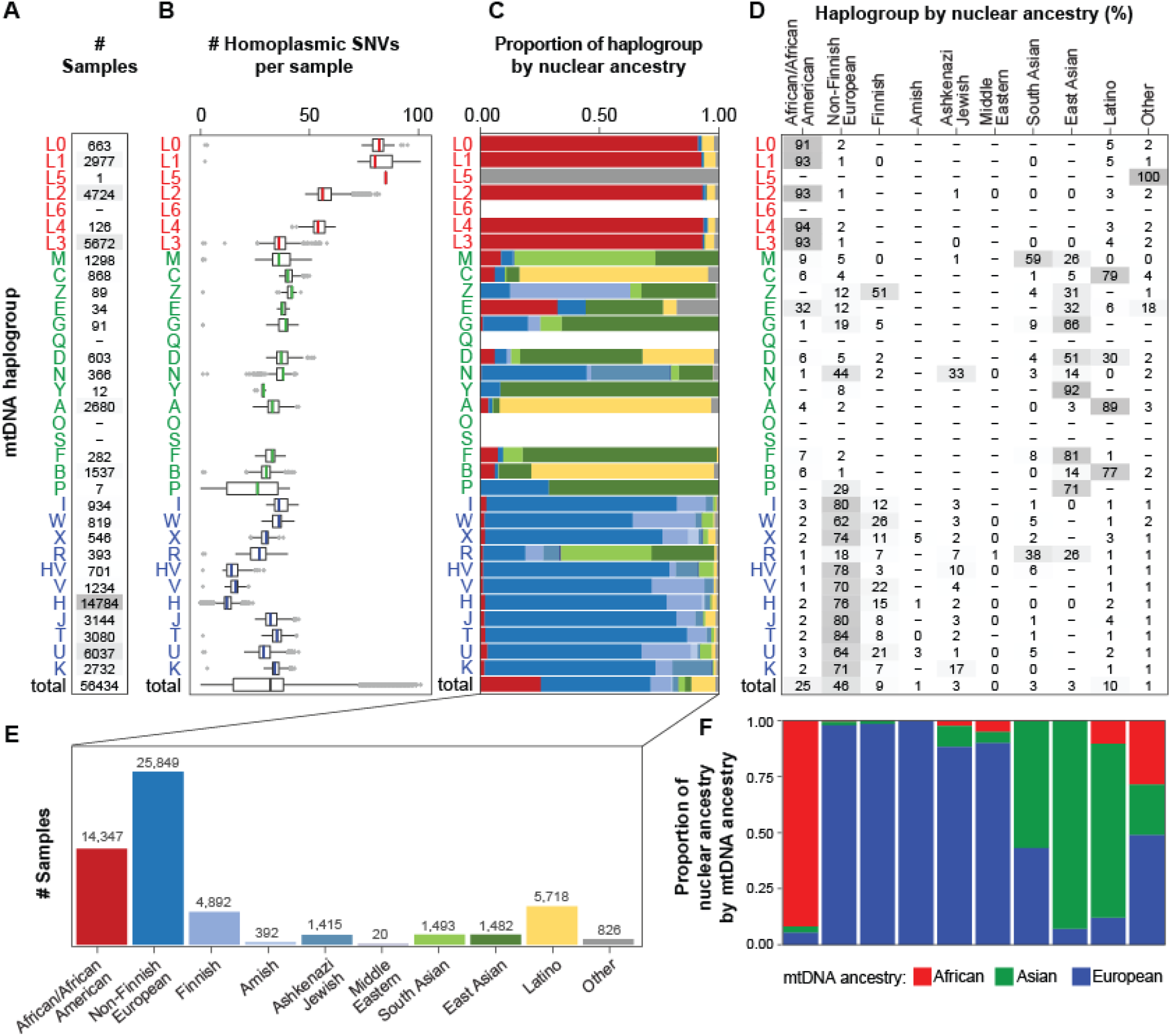
gnomAD v3.1 samples by mtDNA haplogroup and nuclear ancestry. **(A)** The number of samples is shown by mtDNA top-level haplogroup. Color indicates mtDNA haplogroups phylogenetically associated with African (red), Asian (green), or European (blue) origin (Lott et al. 2013). **(B)** For each haplogroup, box plots show the number of homoplasmic SNVs per sample compared to the GRCh38 reference genome (haplogroup H) with median shown in color. **(C)** For each haplogroup, stacked barcharts show nuclear ancestry from nuclear genome analysis, with colors as in panel E. **(D)** For each haplogroup, the percentage of samples from each inferred nuclear ancestry is shown in a heatmap. Dash indicates 0 samples, while 0 indicates a percentage between 0-1. **(E)** The number of samples is shown by inferred nuclear ancestry. **(F)** For each inferred nuclear ancestry shown in panel **D**, stacked barchart shows mtDNA haplogroups phylogenetically associated with African (red), Asian (green), or European (blue) origin (Lott et al. 2013).

GnomAD annotates sample ancestry based on principal components analysis of the nuclear genome (Karczewski et al. 2020) (Fig. 4C–E). The 56,434 samples were predominantly of European (58%) and African (25%) ancestry with lower representation from Latino and admixed American (10%), East Asian (3%), and South Asian (3%) ancestral populations (Fig. 4E). The mtDNA haplogroups were largely concordant with nuclear ancestry (Fig. 4C, F).

### Patterns of variation in mtDNA genes

Unlike the nuclear genome, approximately 90% of the mtDNA encodes protein or RNA genes, and only 10% is intergenic. The proportion of possible SNVs observed was consistent with selection against non-synonymous and RNA variation. Specifically, 55% of all possible synonymous variants were observed, but only 10% of possible missense and RNA variants, and 1% of possible stop gain variants were observed (Fig. 5A). We also observed fewer possible SNVs in the non-coding control region compared to synonymous variants (Fig. 5A), and this held true within the hypervariable region and when limiting to transitions (Fig. S6A). The proportion of variants observed at homoplasmy and the median maximum heteroplasmy of heteroplasmic variants decreased as the predicted severity of the variant type increased (Fig. 5B and Fig. S6B-D). SNV and indel variants in the RNA genes showed a similar pattern of heteroplasmy to each other. Only two predicted loss-of-function variants were homoplasmic in gnomAD (1 stop gain and 1 frameshift). However, manual inspection revealed neither is likely a true loss-of-function, as the frameshift can result in a protein of the same length, and the stop gain is rescued by a multi-nucleotide variant in the same codon (Table S3).

**Figure 5:**
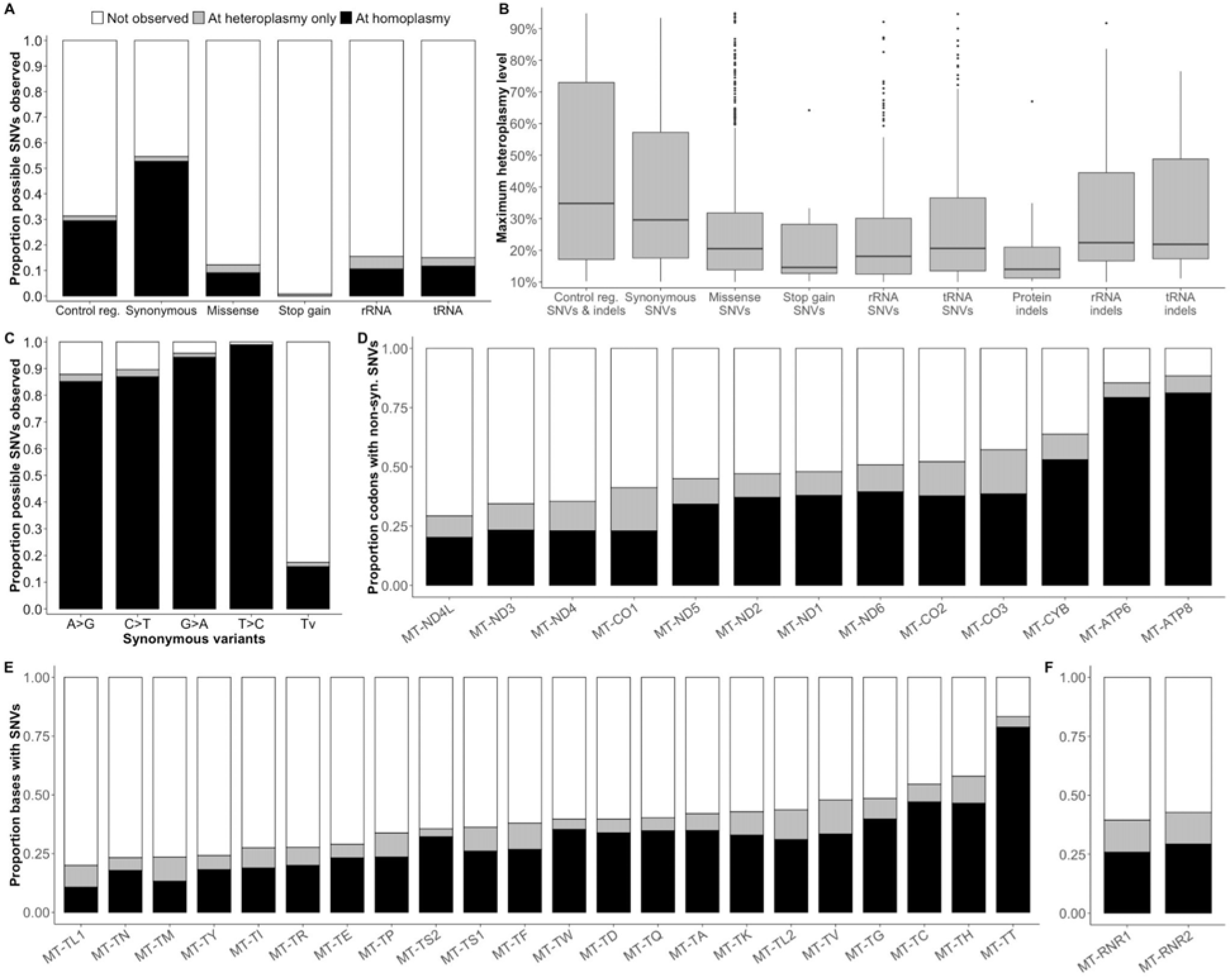
Patterns of variation in the mtDNA in gnomAD. **(A)** The barchart shows the proportion of possible SNVs observed, partitioned into those observed at homoplasmy (black), only at 10-95% heteroplasmy (gray), or not observed (white). **(B)** The boxplot shows the maximum heteroplasmy of variants observed only at heteroplasmy. Protein indels include frameshift and in-frame variants. ‘Control reg.’ represents the non-coding control region m.16024-576 in **(A)** and **(B)**. **(C)** The barchart shows the proportion of possible synonymous variants observed in gnomAD for transversions (Tv) and all possible transitions (A>G, C>T, G>A, T>C) on the reference strand. **(D)** The barchart shows the proportion of codons in protein-coding genes with non-synonymous SNVs observed. **(E)** and **(F)** show the proportion of bases in tRNA and rRNA genes with SNVs. **(C-F)** follow the color legend in (**A**).

Transitions predominate over transversions across the mtDNA, where T>C and G>A mutations are associated with the highest mutability (Ju et al. 2014). Approximately 95% of possible synonymous T>C and G>A variants were observed at homoplasmy (Fig. 5C), suggesting that the size of this dataset is near saturation for this highly mutable, weakly negatively selected variant type. Interestingly, nearly all of the possible G>A synonymous variants not seen at homoplasmy were within AUG codons that were either a start codon (c.3G>A) or the third codon of a gene with an AUA start codon (Table S4). In the mitochondria, AUG and AUA both code for methionine, although modification of the mitochondrial tRNA^Met^ is required to pair with AUA (Van Haute et al. 2017). In the nine genes with AUG start codons, a c.3G>A variant was never observed at homoplasmy in gnomAD, nor in HelixMTdb, and was absent or seen once in MITOMAP (Table S4) (Lott et al. 2013; Bolze et al. 2020). Collectively, these observations suggest selection against AUA at AUG start codons.

To provide insight into gene-level tolerance of variation, we assessed the proportion of non-synonymous codon changes in protein-coding genes and base changes in RNA genes. Among protein-coding genes, the proportion of codons with a non-synonymous variant ranged from 30-90%, suggesting that some proteins are more tolerant of variation (Fig. 5D). For example, complex V genes *MT-ATP8* and *MT-ATP6* showed the highest proportion of codons with non-synonymous variation, while complex I genes had the lowest proportion. Among the RNA genes, the proportion of bases with a variant ranged from 20-85%, indicating that specific RNAs may be more tolerant of variation, especially *MT-TT* (Fig. 5E–F).

### Prevalence of known pathogenic mtDNA variants in gnomAD

We calculated the carrier frequency of the 94 variants listed as “confirmed” pathogenic in MITOMAP, including 56 reported to cause disease at heteroplasmy (typically > 60% heteroplasmy) and 38 reported to cause disease at homoplasmy or both at homoplasmy and heteroplasmy (Lott et al. 2013; Craven et al. 2017). In gnomAD, we observe 26 pathogenic variants in 231/56,434 individuals, equating to a total carrier frequency of ~1 in 250 individuals (Fig. 6). Fewer variants associated with disease only at heteroplasmy were observed in gnomAD relative to those associated with disease at homoplasmy (16% vs 45%), consistent with the expectation that the latter group includes milder mutations (Fig. S7A) (Craven et al. 2017). Eleven pathogenic variants were observed at homoplasmy, most of which are reported to be incompletely penetrant and/or associated with adult-onset disease when homoplasmic (including nonsyndromic hearing loss, aminoglycoside-induced hearing loss, Leber Hereditary Optic Neuropathy (LHON), or reversible myopathy). One of these variants seen at homoplasmy in gnomAD was not associated with disease at homoplasmy in MITOMAP (m.8993T>C); however, it has recently been described in adult-onset cases at homoplasmy <u>(Stendel et al. 2020)</u>. Across all pathogenic variants, m.1555A>G had the highest carrier frequency (1 in ~750; Fig. 6), while m.3243A>G and m.8344A>G variants had the highest carrier frequency among those only observed at heteroplasmy (~1 in 10,000; Fig. 6).

**Figure 6:**
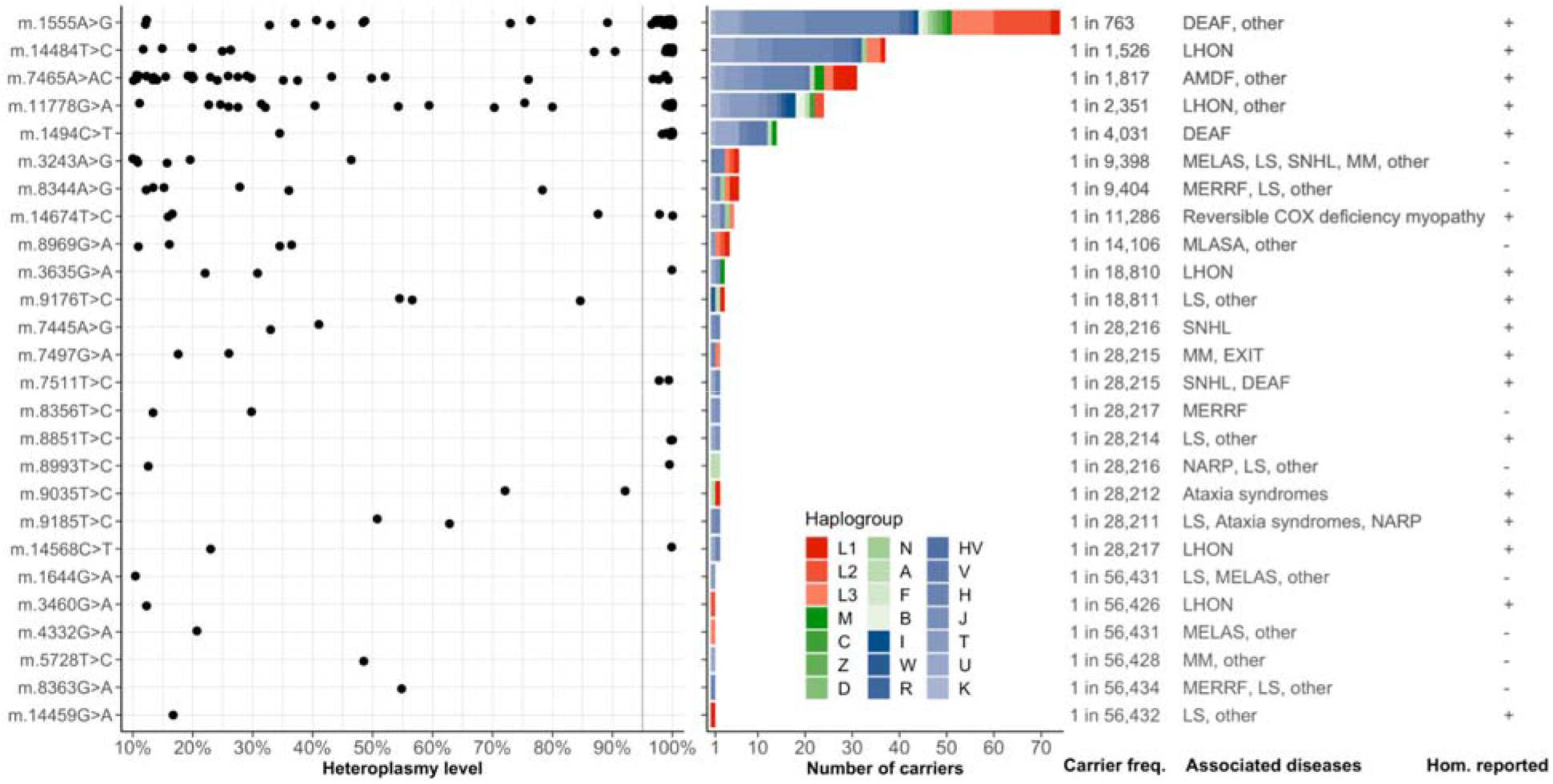
Known pathogenic variants in gnomAD. Shown are the 26 pathogenic variants observed in gnomAD along with their heteroplasmy levels, haplogroup distribution, carrier frequency, MITOMAP-curated disease phenotypes, and indicator showing whether disease occurs at homoplasmy (Hom. reported; note this includes variants only associated with disease at homoplasmy, or at both homoplasmy and heteroplasmy). The carrier frequency is calculated as the high-quality allele count divided by the number of individuals with high quality sequence at the position. The dark grey line at 95% heteroplasmy level represents the threshold used to define homoplasmic variant calls. Haplogroups are ordered by their position in the phylogenetic tree, and colored by their association with African (red), Asian (green), or European (blue) ancestry. AMDF: Ataxia, Myoclonus and Deafness; COX: Cytochrome c Oxidase; DEAF: Maternally inherited deafness or aminoglycoside-induced deafness; EXIT: Exercise Intolerance; LHON: Leber Hereditary Optic Neuropathy; LS: Leigh syndrome; MELAS: Mitochondrial Encephalomyopathy, Lactic Acidosis, and Stroke-like episodes; MERRF: Myoclonic Epilepsy and Ragged Red Muscle Fibers; MLASA: Mitochondrial Myopathy, lactic acidosis and sideroblastic anemia; MM: Mitochondrial Myopathy; NARP: Neurogenic muscle weakness, Ataxia, and Retinitis Pigmentosa; SNHL: Sensorineural Hearing Loss; other: other phenotypes listed for this variant in MITOMAP.

Mitochondrial DNA specifications of the American College of Medical Genetics and Association of Molecular Pathology (ACMG/AMP) guidelines for sequence variant interpretation state that allele frequency in population databases < 0.00002 or > 0.005 can provide evidence of pathogenicity or benign impact respectively, where analysis of homoplasmic databases was used to determine these thresholds (McCormick et al. 2020). Consistent with this guideline, none of the pathogenic mtDNA variants in gnomAD had a homoplasmic allele frequency (AF_hom) that satisfied benign variant frequency criteria (AF_hom > 0.005 for benign strong BS1, or AF_hom > 0.01 for benign stand-alone BA1). Approximately 90% of the 94 known pathogenic variants had AF_hom < 0.00002, satisfying the pathogenic supporting criteria PM2_supporting for variant frequency; this included all of the variants only associated with disease at heteroplasmy (Fig. S7B). All pathogenic variants also had AF_hom < 0.005 within haplogroups and populations (Fig. S7B). Analysis of the heteroplasmic allele frequency (AF_het) of pathogenic variants showed that all were < 0.005, and that ~85% were < 0.00002 (Fig. S7B). Consistent with recent observations in the UK Biobank and HelixMTdb, the AF_hom of m.14484T>C was greater than the maximum credible population AF reported by Bolze *et al* (0.00053 vs 0.00023), lending support to the suggestion that this variant alone may not cause LHON (Bolze et al. 2020).

## DISCUSSION

Here we present a pipeline for calling homoplasmic and heteroplasmic mtDNA variants and its application to gnomAD v3.1. To our knowledge, this represents the first database of mtDNA variants from WGS data and the only database with heteroplasmic variants aside from HelixMTdb. We present a conservative set of variants based on WGS from 56,434 individuals, after stringent filtering of samples with low mtDNA copy number, samples derived from cell lines, and samples with high contamination. Moreover, we have chosen to report heteroplasmic variants that are occurring at a level of 10% or greater. As expected, the vast majority of variant calls were homoplasmic, including nearly all known haplogroup markers and thousands of additional rare homoplasmic variants. Most heteroplasmies occurred at variants that were observed at homoplasmy in at least one individual. The gnomAD dataset and web browser provides detailed information for each variant, including predicted functional consequence, distribution of heteroplasmy levels, maximum observed heteroplasmy, and population allele frequencies (including aggregated per haplogroup and per nuclear ancestry population).

Our analyses show that misalignment of polymorphic NUMTs contributes to alarmingly high false positive mtDNA variant calls in WGS, particularly for variants with low putative heteroplasmy and for samples with low mtDNA copy number (mtCN) (Fig. 2D,K). Using two polymorphic NUMTs not in the reference human genome assembly that we validated using PacBio sequencing, we estimate a lower bound for NUMT-derived false positives (NUMT-FPs). In samples with mtCN < 50, NUMT-FPs account for the majority of putative heteroplasmies VAF 0.01-0.10 (Fig. 2F). Conversely, in samples with mtCN > 500 (e.g., tissues and cell lines), NUMT-FPs show putative heteroplasmy substantially less than 0.01, and thus are typically not a problem. Even after excluding samples with mtCN < 50 and mtCN > 500, we observe that 50% of variants with VAF 0.01 are NUMT-FPs (Fig. 2K). These NUMT-FPs are also called by tools such as mToolBox that exclude reads mapping to both mitochondrial and nuclear genomes, since such approaches cannot account for polymorphic NUMTs. Our data suggest that for Illumina WGS data, with insert sizes ~375bp, the NUMTs in the reference genome do not cause substantial false positives whereas reads from NUMTs that are not found in the reference genome will misalign to the mtDNA genome. To our knowledge, there are no estimates of NUMT-FPs from HelixMTdb or WGS studies focusing on heteroplasmies (Wei et al. 2019).

Given these findings, and to avoid NUMT-FPs, we employ stringent sample filtering and release only variants with heteroplasmy ≥ 10%. Future releases may develop more sophisticated approaches to define a sample-specific heteroplasmy threshold to exclude NUMT-derived artifacts or may reduce the threshold to 5%. Ultimately, long-read sequencing technologies will be required to fully address the NUMT misalignment problem.

Our data show that known cell lines harbor excess heteroplasmies, including excess deleterious variants (Fig. 2H). These findings likely result from relaxed selection pressures in high-glucose cell culture conditions that are tolerant to ordinarily deleterious mtDNA variants. We note that this finding is of particular importance given emerging technologies that culture patient-derived cells *ex vivo* before transplantation into the individuals (e.g. CAR-T and stem cell therapies), and may warrant further study (Perales-Clemente et al. 2016).

Our pipeline has several limitations. The pipeline is available and easy to run within a scalable cloud-based framework but may be difficult to implement on local compute resources. However, the Mutect2 “mitochondrial-mode” variant caller is easy to run and provides comparable results to other stand-alone tools such as mtDNA-Server and mToolBox. These tools show different trade-offs; e.g. mtDNA-Server shows higher sensitivity for variants with 1% heteroplasmy at the cost of reduced precision. For analysis of other cohorts, less stringent sample and heteroplasmy filtering may be more appropriate depending on mtCN observed in the cohorts.

Analyses of variants in gnomAD are broadly consistent with previous studies of human mtDNA variation. We observed a carrier frequency of ~1 in 250 individuals (VAF 0.10-1.00), consistent with estimates from other studies (Elliott et al. 2008; Wei et al. 2019). Our observed patterns of variation suggest negative selection against variants which impair gene function, as reported by others (Stewart et al. 2008; Wei et al. 2019; Bolze et al. 2020). Missense, tRNA, and rRNA variants showed similar occurrence and heteroplasmy distributions, suggesting they may be removed from the population at a similar rate by negative selection. We observed less variation within the non-coding control region compared to synonymous variants at transition variants (but not at transversion variants; Fig. S6A); however, this may be explained by the higher prevalence of the most mutable tri-nucleotides at synonymous sites (Zhou et al. 2014). To our knowledge, our analyses are the first to reveal a lack of putative synonymous variants at start codons, changing AUG>AUA, implying such mutations may impair mitochondrial function and fitness. Studies in bacteria and yeast mitochondria have shown that AUG is a more efficient initiation codon than AUA (Romero and García 1991; Mulero and Fox 1994). The identification of a c.3G>A variant in an individual with mitochondrial disease may thus warrant further investigation.

We anticipate that gnomAD mtDNA variants will be of broad use in the clinical interpretation of variants; however, we want to emphasize key limitations for interpretation of heteroplasmic variants detected in patients. The mitochondrial specifications of the ACMG/AMP guidelines provides clear methods for variant interpretation based on homoplasmic allele frequency (AF_hom): specifically, AF_hom > 0.005 provides evidence for benign classification whereas AF_hom < 0.00002 is supporting evidence for pathogenicity (McCormick et al. 2020). However, for variants never detected at homoplasmy, no such guidelines for heteroplasmic allele frequency (AF_het) have yet been developed. In the default browser setting, we have chosen to only include variants that we observe at heteroplasmy ≥ 10%, because below this threshold we observed thousands of variants that are enriched for NUMT-derived false positives and sequencing errors. It is important to note that if clinical sequencing of a patient detects a low heteroplasmy variant (e.g., heteroplasmy less than 10-15%) that is apparently absent from gnomAD based on the browser view, we caution against using the absence to support pathogenicity, and urge gnomAD users to select the “Include unfiltered variants” option to view these artifact-prone sites and other excluded variants. These filtered variants are also included in downloadable gnomAD data files with the relevant flags. This scenario applies specifically to low heteroplasmy variants -- which are prone to sequencing errors and NUMT-misalignment that are not typically problematic for high heteroplasmy mtDNA variants or nuclear variants.

Given the challenge and extent of NUMT-derived false positives, we urge confirmatory studies of putative low level heteroplasmy variants detected by clinical diagnostics. We note that many clinical sequencing methods (appropriately) aim to avoid NUMT-derived artifacts, using specialized methods to enrich for circular DNA or long-range PCR that selectively amplifies intact mtDNA. However, even such specialized methods may inadvertently report NUMT-derived false positives, as may be the case in the controversial report of paternally inherited mtDNA (Luo et al. 2018; Lutz-Bonengel and Parson 2019; Lutz-Bonengel et al. 2021).

GnomAD’s diverse population representation, exclusion of individuals known to have severe pediatric disease, and capture of homoplasmic and heteroplasmic variation offer value for mtDNA variant interpretation. As the first large-scale mtDNA database built from WGS data via a publicly available pipeline, this study has provided both open-source tools and data that will support mtDNA analysis as part of clinical WGS testing.

## METHODS

### Mitochondrial variant calling pipeline in single samples

Whole genome sequence data were aligned to reference genome GRCh38, which includes chrM (identical to the revised Cambridge Reference Sequence, GenBank NC_012920.1) using BWAmem version 0.7.15-r1140 (parameters -K 100000000 -p -v 3 -t 2 -Y). For each sample CRAM, Terra MitochondrialPipeline version 25 was run (https://portal.firecloud.org/?return=terra#methods/mitochondria/MitochondriaPipeline/25). Briefly, GATK version 4.1.2.0 (McKenna et al. 2010) tools were used to estimate the median nuclear genome coverage (Picard CollectWgsMetrics), to exclude duplicates (Picard MarkDuplicates), to pull reads from chrM (GATK PrintReads --read-filter MateOnSameContigOrNoMappedMateReadFilter --read-filter MateUnmappedAndUnmappedReadFilter), and to call variants (GATK Mutect2 --mitochondria-mode -- annotation StrandBiasBySample --max-reads-per-alignment-start 75 --max-mnp-distance 0). For calling variants in the control region (coordinates chrM:16024-16569 and chrM:1-576), reads originally aligning to chrM were realigned to a chrM reference genome shifted by 8000 nucleotides, and then variants called on the shifted reference were mapped back to standard coordinates (Picard LiftOver) and combined with variants from the non-control region. Mutect2 variants were then filtered (GATK FilterMutectCalls --stats raw_vcf_stats --max-alt-allele-count 4 --mitochondria-mode --autosomal_coverage nDNA_MEDIAN_COV --min_allele_fraction 0.01); multi-allelic sites were split into different variants (LeftAlignAndTrimVariants --split-multi-allelics --dont-trim-alleles --keep-original-ac); and HaploGrep/HaploCheck (v1.0.5) was run to assign haplogroup (Weissensteiner et al. 2016b) and estimate mtDNA contamination (Weissensteiner et al. 2021). The min_vaf_threshold was set to 0.01 and calls below 0.01 VAF were later set to homoplasmic reference. For each input sample, a VCF with mtDNA variants was produced. We note that GATK left-aligns all indel calls, unlike calls from mtDNA-Server and variants in the Phylotree database.

We developed Mutect2 “mitochondria mode,” which, in contrast to its original use in calling somatic mutations in cancer, sets parameters and filters specialized for calling low VAF variants in high coverage mtDNA. Mutect2 performs local read realignment (using the same realignment algorithm as GATK’s HaplotypeCaller), performs a local assembly of haplotypes, prunes these haplotypes, and then calls SNVs and short insertions/deletions via a Bayesian somatic genotyping model. To increase sensitivity, Mutect2 “mitochondria mode” lowers the threshold for ActiveRegions (regions to be considered by the variant caller) and the threshold for emitting variants based on quality. Additionally, “mitochondria mode” implements a specialized adaptive approach to prune paths from the assembly graph, which is necessary due to extremely high mtDNA coverage. Adaptive pruning uses both the local coverage and observed sequencing error rate to determine appropriate paths to prune from the graph to reduce false positive calls. Finally, “mitochondria mode” removes several standard Mutect2 filters (including clustered events, filtered haplotypes, and multiallelic) that operate with the assumption that variants do not typically occur near each other, which does not apply to mtDNA.

A predefined list of artifact-prone sites (positions 301, 302, 310, 316, 3107, 16182) was provided as input into this pipeline, and any variant overlapping these sites was filtered (“artifact_prone_site”), similar to other tools (Weissensteiner et al. 2016a; Wei et al. 2019). Sequence context at these specific artifact-prone sites makes it difficult to distinguish true variants from technical artifacts. The homopolymer tracts at location chrM:300-317 (AAACCCCCCCTCCCCCGC) cause Illumina sequencing errors in all samples and cause (i) a large coverage dip in this region, (ii) reads with many apparent indels near position chrM:310T, and (iii) apparent substitutions of chrM:301A>C, chrM:302A>C, chrM:310T>C, and chrM:316G>C. Similarly, homopolymer tracts at location chrM:16180-16193 (AAAACCCCCTCCCC) cause errors and apparent indels at position chrM:16182-16183. The reference genome contains “N” at position chrM:3107, which causes misalignment of many reads. We note that this artifact-prone site filter was re-implemented at the cohort level after variants were combined across samples (see below).

### Reproducibility

Duplicates were determined as described in Karczewski *et al* (2020). For Mutect2, we ran version 25 of the Terra MitochondrialPipeline, filtered artifact-prone sites, and set any filtered genotypes to homoplasmic reference. To measure how similar variant calls were between duplicate samples for each tool, we calculated the jaccard index for all variants as well as only variants with VAF > 0.10, 0.50, and 0.90. We output the results of this comparison for both SNVs and indels, but note that mtDNA-Server is focused on calling SNVs and that their method for calling indels is in beta testing.

### Assessing accuracy on truth datasets

Sample NA12878 and 22 samples from diverse L haplogroups were selected for in silico mixing experiments to create a large truth dataset compared to the reference chrM (totalling 1,200 variants at 286 positions, including 8 indels). For each L-haplogroup sample, the number of mtDNA reads per sample was counted (samtools v1.8 idxstats), and then downsampling was performed (samtools v1.8) to create five BAM files containing a predefined ratio of reads from the L-haplogroup sample and NA12878 (0%, 1%, 5%, 90%, 99%, 100%). For each mixture, total coverage was set to the L-haplogroup sample’s original coverage. GATK’s HaplotypeCaller version 4.0.3.0 was used to call homoplasmic variants on the original BAMs before downsampling, with the ploidy argument set to 100. For each L haplogroup sample, a truth set was defined as variants present in the L haplogroup sample (allele count > 94/100) but absent in NA12878 (based on manual review using over-lapping read pair data, with padding of 1bp around each NA12878 variant). For each L-haplogroup sample mixture, true and false positive calls were calculated against the sample-specific truth set, and then summed across all 22 L-haplogroup samples to create sensitivity and precision metrics.

### mtDNA-Server comparison

We used Hail’s Batch service (Hail Team) to run mtDNA-Server on sample mixtures and sample duplicates. We ran mutserve v1.3.4 using “analyse-local” with the heteroplasmy level threshold set to 0.01 and parameters outputting deletions and insertions. Consecutive deletion calls were merged (if VAF differed < 0.10) and summarized with mean depth and heteroplasmy. Output was reformatted to match Mutect2 calls. Bcftools (v1.10.2) (Li 2011) was used to left-align and normalize the variants.

### NUMT-derived false positives

We identified candidate NUMT-FPs (Table S1) as follows: we identified 122 common heteroplasmic mtDNA variants (unfiltered variants with VAF 0-0.50 in ≥ 1000 samples), of which 67 had heteroplasmy levels that correlated with 1/(1+mtCN) (Spearman correlation > 0.45), of which over half co-occurred with another common heteroplasmy in the same samples (Pearson correlation > 0.45). For each candidate NUMT-FP, we generated a 20mer sequence centered on the variant, then searched the derived 20mer (and its reverse complement) against PacBio SAM files corresponding to three cell lines (NA12891, NA19239, NA19238), with read lengths ~10Kb. PacBio reads that contained the 20mers were aligned to GRCh38 via Blat (Kent 2002). For two NUMTs, exact NUMT sequence and break points were identified that give rise to 25 validated NUMT-FPs (Table S1). We defined “linked NUMT-FPs” as those where at least two of the 25 NUMT-FPs derived from the same NUMT were present in the same sample with heteroplasmy levels within 0.05 (unfiltered variants, VAF 0-0.50). For Fig. 2F, all PASS variants were binned by VAF (e.g. 0.01-0.02); all samples were binned by sample mtCN (e.g. 25-50); and then the fraction of all variants in those bins that were “linked NUMT-FPs” was calculated and plotted.

### PacBio sequencing and data processing

We performed long read sequencing using the Pacific Biosciences (PacBio) circular consensus sequencing (CCS) protocol. Briefly, for library preparation, we obtained 5 μg of high molecular weight genomic DNA (> 50% of fragments ≥ 40 kb) and sheared fragments to ~10 kb using the Megaruptor 3 (B06010003; Diagenode), followed by DNA repair and ligation of PacBio adapters using the SMRTbell Template Prep Kit v1.0 (100-991-900). Each library was subsequently size selected for 10 kb ± 20% using the SageELF with 0.75% agarose cassettes (Sage Science). Libraries were quantified with the Qubit dsDNA High Sensitivity Assay Kit (Q32854; Thermo Fisher Scientific), subsequently diluted to 50 pM per single molecule, real-time (SMRT) cell, hybridized with PacBio v2 sequencing primer, and bound with SMRT sequencing polymerase using Sequel II Binding Kit 1.0 (101-731-100). Sequencing was performed in CCS mode on the Sequel II instrument using 8M SMRT Cells (101-389-001) and Sequel II Sequencing 1.0 Kit (101-717-200), with a 2-hour pre-extension time and 30-hour movie time per SMRT cell. Initial quality filtering, base calling, and adapter marking were performed automatically on-board the Sequel II to generate an initial raw “subreads.bam” file. CCS reads were generated using CCS software v.3.4.1 from PacBio (https://github.com/PacificBiosciences/ccs) with parameters “--minPasses 3 -- minPredictedAccuracy 0.99 --maxLength 21000.” CCS reads were mapped to the “GRCh38_noalt” reference sequence (GRCh38 without decoy sequences, HLA sequences, and alternative loci representations) using minimap2 (version 2.17-r941 with parameters “-ayYL --MD --eqx -x map-pb”).

### Cell line analyses

Selected gnomAD cohorts were annotated as cell lines including samples from 1000 Genomes Project and the Human Genome Diversity Project (n=3,277), and Osaka University (n=246). Variant subtypes for known cell lines and samples with mtCN50-500 were annotated via the Variant Effect Predictor (VEP) and categorized as pLOF (if VEP consequence=stop_gained|frameshift_variant), missense (VEP consequence=missense_variant), synonymous (VEP impact=LOW), or rRNA/tRNA (VEP biotype=Mt_tRNA|Mt_rRNA); otherwise they were categorized as non-coding.

### gnomAD sample and variant filtering

gnomAD v3.1 contains 76,156 samples passing filters, of which 70,375 had read data available for analysis. For mtDNA analysis we analyzed 56,434 samples after excluding 6,505 samples with mtCN < 50, 5633 samples with mtCN > 500, and 1803 samples with contamination > 2% based on nuclear contamination (VerifyBamID v1, v2) (Jun et al. 2012; Zhang et al. 2020), mtDNA contamination (Haplocheck v1.0.5) (Weissensteiner et al. 2021), or an internal algorithm (mt-high-hets). Mt-high-hets utilizes the PASS haplogroup-defining variants which should be homoplasmic (VAF=1.00), but in contaminated samples show multiple alleles with VAF 0.85-0.998. Mt-high-hets calculates contamination = 1-mean(VAF 0.85-0.998) if 3 such variants are present; otherwise contamination = 1-mean(VAF 0.85-1.00).

To distinguish between missing calls and homoplasmic reference sites after combining the samples, we set the genotype of a sample that lacked a call at a site to homoplasmic reference if the depth of coverage at the respective site was greater than 100x. The genotype was otherwise set to missing.

Problematic variants were filtered or flagged as follows. Flag “common_low_heteroplasmy” was applied to variants found with PASS genotypes in > 56 individuals (allele frequency > 0.001) at VAF 0-0.50. (Note this includes PASS variants 0-0.01 VAF, which are subsequently filtered.) Filter “indel_stack” was applied to any indel allele where all samples with a variant call had at least 2 different heteroplasmic indels called at that position. The Hail pipeline re-implemented the “artifact_prone_sites” filter and any variant overlapping positions 301, 302, 310, 316, 3107, or 16182 was filtered. The original single-sample pipeline assigned filters “possible_numt” and “mt_many_low_hets” which were found to be unreliable and were ignored in the gnomAD release. Filter “npg” (no pass genotype) was applied to variants which had no passing genotype across all samples.

### gnomAD annotations

All variant annotations were implemented in Hail. Annotations from VEP (v101) (McLaren et al. 2016) were added using the same pipeline which was used for gnomAD v3.1 nuclear annotations, with the modification of changing the distance parameter to 0 to avoid “upstream” and “downstream” annotations. We obtained rsIDs from dbSNP (b154) and added in silico prediction annotations for tRNA variants from PON-mt-tRNA (download date 2020-08-27) (Niroula and Vihinen 2016), MitoTIP (download date 2020-08-27) (Sonney et al. 2017), and HmtVar (Preste et al. 2019). We define heteroplasmic variants as variants with VAF 0.10-0.95 and homoplasmic variants as variants with VAF 0.95-1.00. We generated allele frequency information for both heteroplasmic and heteroplasmic variants and also provided this information for each top-level haplogroup and population.

### Multi-nucleotide variants (MNVs)

Homoplasmic MNVs that were found adjacent in at least 90% of samples were flagged on the web portal. Specifically, homoplasmic MNVs were defined as variants where AC_hom_MNV/AC_hom > 0.90, where AC_hom_MNV indicates the number of samples where this homoplasmic variant was adjacent to any other homoplasmic variant, and AC_hom indicates the number of samples with this homoplasmic variant. For example, adjacent variants chrM:5185G>A (homoplasmic in 1 sample) and chrM:5186A>T (homoplasmic in 80 samples) were observed together in 1 sample and thus the former was flagged MNV (AC_hom_MNV/AC_hom = 1/1) whereas the latter was not (AC_hom_MNV/AC_hom = 1/80).

### Haplogroups

Haplogroups were downloaded from the rCRS-orientated version of PhyloTree Build 17 (van Oven and Kayser 2009), and variants were extracted using custom Python scripts. As Phylotree represents a right-alignment of indels, we manually inspected haplogroup indel variants and inferred the equivalent left-alignment that would be expected in gnomAD, with the exception of haplogroup insertions of unknown length (denoted by ‘.X’).

### Inferred nuclear ancestry

Each sample was assigned to a predefined set of ancestries (Fig. 4E), based on principal component analysis of nuclear SNVs (Karczewski et al. 2020)

### Proportion possible observed

A ‘synthetic’ VCF with all possible mtDNA SNVs was generated using an in-house script, and annotated by VEP (v97). This was used to calculate the proportion of possible SNVs observed in gnomAD. For variants within two genes (with two consequences), both consequences were included in the possible variant counts. The proportion of codons in each protein-coding gene with homoplasmic or only heteroplasmic non-synonymous SNVs (all SNVs except those with consequence “synonymous_variant”), and the proportion of bases in each RNA gene with homoplasmic or only heteroplasmic SNVs in gnomAD were calculated using a custom script.

### Pathogenic variants and other variant annotations

Pathogenic variants with a ‘Confirmed’ status were downloaded from the MITOMAP database (download date 2021-02-22) (Lott et al. 2013); indel variants were manually inspected and the equivalent left-alignment that would be expected in gnomAD was inferred. APOGEE in silico predictions were downloaded from MitImpact (v3.0.6) (Castellana et al. 2017). HmtVar in silico predictions were retrieved from the HmtVar database (download date 2020-11-18) (Preste et al. 2019). Maximum heteroplasmy data from HelixMTdb was downloaded from Helix.com (version dated 03/27/2020) (Bolze et al. 2020).

### DATA ACCESS

Variants and population frequencies are available in gnomAD v3.1 (gnomad.broadinstitute.org). A user-friendly website provides variant annotations, including distributions across heteroplasmy levels, populations, and haplogroups. Data are available for download in multiple formats, including VCF, Hail Table, and simple tab-delimited files (https://gnomad.broadinstitute.org/downloads#v3-mitochondrial-dna). The Mutect2 pipeline is available through GATK at https://github.com/broadinstitute/gatk/blob/master/scripts/mitochondria_m2_wdl/MitochondriaPipeline.wdl (the data available in gnomAD v3.1 was generated using https://portal.firecloud.org/?return=terra#methods/mitochondria/MitochondriaPipeline/25), and the Hail scripts used for combining the VCFs, filtering samples and variants, adding annotations, and performing analyses can be found at https://github.com/broadinstitute/gnomad-mitochondria.

## ACKNOWLEDGMENTS

We thank David Thorburn, John Christodoulou, and members of their labs for their feedback on displaying mtDNA variants on the browser. The gnomAD results published here are in part based on data: (1) generated by The Cancer Genome Atlas (TCGA) managed by the NCI and NHGRI (accession: phs000178.v10.p8); information about TCGA can be found at http://cancergenome.nih.gov; (2) generated by the Genotype-Tissue Expression Project (GTEx) managed by the NIH Common Fund and NHGRI (accession: phs000424.v7.p2); (3) generated by the Alzheimer’s Disease Sequencing Project (ADSP), managed by the NIA and NHGRI (accession: phs000572.v7.p4). Analysis of the Genome Aggregation Database was supported by NIDDK U54DK105566 and the National Human Genome Research Institute of the National Institutes of Health under award number U24HG011450. N.J.L. received a National Health and Medical Research Council (NHMRC) Early Career Fellowship and an Australian American Association Scholarship. This work was supported by grants from the Broad Institute Scientific Projects to Accelerate Research and Collaboration (S.E.C and V.K.M) grant and the National Institutes of Health (R35GM122455 (S.E.C. and V.K.M). Additional funding for Genome Aggregation Database Consortium members is listed in Supplemental Materials. The content is solely the responsibility of the authors and does not necessarily represent the official views of the National Institutes of Health.

## Author Contributions

K.M.L and N.J.L performed data analysis and manuscript preparation. N.A.W. adapted the gnomAD browser to incorporate mitochondria-specific annotations. M.S. and D.B. developed the mitochondria mode of Mutect2. M.S., A.H, and J.S. developed the mitochondrial pipeline and Terra WDL with guidance from L.G., E.B., and assistance from J.E. K.G. developed the PacBio sequencing pipeline. GnomAD Consortium provided samples and project oversight. H.L.R and D.G.M provided gnomAD leadership. G.T. provided oversight of the collaboration and manuscript preparation. M.L., and V.K.M. provided leadership, analytical advice, and editing of the manuscript. S.E.C. provided leadership, expert knowledge, data analysis, and manuscript preparation. All authors listed under The Genome Aggregation Database Consortium contributed to the generation of the primary data incorporated into the gnomAD resource and specific members provided additional contributions: Mark Daly provided gnomAD leadership, Sebastian Schönherr provided assistance with Haplocheck, Konrad Karczewski reviewed the code for call set assembly and annotation.

## REFERENCES

Anderson S, Bankier AT, Barrell BG, de Bruijn MH, Coulson AR, Drouin J, Eperon IC, Nierlich DP, Roe BA, Sanger F, et al. 1981. Sequence and organization of the human mitochondrial genome. Nature 290: 457–465.

Andrews RM, Kubacka I, Chinnery PF, Lightowlers RN, Turnbull DM, Howell N. 1999. Reanalysis and revision of the Cambridge reference sequence for human mitochondrial DNA. Nat Genet 23: 147.

Benjamin DI, Sato T, Cibulskis K, Getz G, Stewart C, Lichtenstein L. 2019. Calling Somatic SNVs and Indels with Mutect2. BioRxiv doi: 10.1101/861054.

Bolze A, Mendez F, White S, Tanudjaja F, Isaksson M, Jiang R, Rossi AD, Cirulli ET, Rashkin M, Metcalf WJ, et al. 2020. A catalog of homoplasmic and heteroplasmic mitochondrial DNA variants in humans. BioRxiv doi: 10.1101/798264.

Brown WM, George M, Wilson AC. 1979. Rapid evolution of animal mitochondrial DNA. Proc Natl Acad Sci USA 76: 1967–1971.

Calabrese C, Simone D, Diroma MA, Santorsola M, Guttà C, Gasparre G, Picardi E, Pesole G, Attimonelli M. 2014. MToolBox: a highly automated pipeline for heteroplasmy annotation and prioritization analysis of human mitochondrial variants in high-throughput sequencing. Bioinformatics 30: 3115–3117.

Cann RL, Stoneking M, Wilson AC. 1987. Mitochondrial DNA and human evolution. Nature 325: 31–36.

Castellana S, Fusilli C, Mazzoccoli G, Biagini T, Capocefalo D, Carella M, Vescovi AL, Mazza T. 2017. High-confidence assessment of functional impact of human mitochondrial non-synonymous genome variations by APOGEE. PLoS Comput Biol 13: e1005628.

Cavalli-Sforza LL. 1998. The DNA revolution in population genetics. Trends Genet 14: 60–65.

Clima R, Preste R, Calabrese C, Diroma MA, Santorsola M, Scioscia G, Simone D, Shen L, Gasparre G, Attimonelli M. 2017. HmtDB 2016: data update, a better performing query system and human mitochondrial DNA haplogroup predictor. Nucleic Acids Res 45: D698–D706.

Craven L, Alston CL, Taylor RW, Turnbull DM. 2017. Recent advances in mitochondrial disease. Annu Rev Genomics Hum Genet 18: 257–275.

Dayama G, Emery SB, Kidd JM, Mills RE. 2014. The genomic landscape of polymorphic human nuclear mitochondrial insertions. Nucleic Acids Res 42: 12640–12649.

Elliott HR, Samuels DC, Eden JA, Relton CL, Chinnery PF. 2008. Pathogenic mitochondrial DNA mutations are common in the general population. Am J Hum Genet 83: 254–260.

Gorman GS, Chinnery PF, DiMauro S, Hirano M, Koga Y, McFarland R, Suomalainen A, Thorburn DR, Zeviani M, Turnbull DM. 2016. Mitochondrial diseases. Nat Rev Dis Primers 2: 16080.

Gorman GS, Schaefer AM, Ng Y, Gomez N, Blakely EL, Alston CL, Feeney C, Horvath R, Yu-Wai-Man P, Chinnery PF, et al. 2015. Prevalence of nuclear and mitochondrial DNA mutations related to adult mitochondrial disease. Ann Neurol 77: 753–759.

Grady JP, Pickett SJ, Ng YS, Alston CL, Blakely EL, Hardy SA, Feeney CL, Bright AA, Schaefer AM, Gorman GS, et al. 2018. mtDNA heteroplasmy level and copy number indicate disease burden in m.3243A>G mitochondrial disease. EMBO Mol Med 10: e8262.

Hail Team. Hail 0.2. https://github.com/hail-is/hail

Jun G, Flickinger M, Hetrick KN, Romm JM, Doheny KF, Abecasis GR, Boehnke M, Kang HM. 2012. Detecting and estimating contamination of human DNA samples in sequencing and array-based genotype data. Am J Hum Genet 91: 839–848.

Ju YS, Alexandrov LB, Gerstung M, Martincorena I, Nik-Zainal S, Ramakrishna M, Davies HR, Papaemmanuil E, Gundem G, Shlien A, et al. 2014. Origins and functional consequences of somatic mitochondrial DNA mutations in human cancer. elife 3: e02935.

Karczewski KJ, Francioli LC, Tiao G, Cummings BB, Alföldi J, Wang Q, Collins RL, Laricchia KM, Ganna A, Birnbaum DP, et al. 2020. The mutational constraint spectrum quantified from variation in 141,456 humans. Nature 581: 434–443.

Kent WJ. 2002. BLAT--the BLAST-like alignment tool. Genome Res 12: 656–664.

Lek M, Karczewski KJ, Minikel EV, Samocha KE, Banks E, Fennell T, O’Donnell-Luria AH, Ware JS, Hill AJ, Cummings BB, et al. 2016. Analysis of protein-coding genetic variation in 60,706 humans. Nature 536: 285–291.

Li H. 2013. Aligning sequence reads, clone sequences and assembly contigs with BWA-MEM. arXiv:13033997v2.

Li H. 2011. A statistical framework for SNP calling, mutation discovery, association mapping and population genetical parameter estimation from sequencing data. Bioinformatics 27: 2987–2993.

Li M, Schroeder R, Ko A, Stoneking M. 2012. Fidelity of capture-enrichment for mtDNA genome sequencing: influence of NUMTs. Nucleic Acids Res 40: e137.

Lott MT, Leipzig JN, Derbeneva O, Xie HM, Chalkia D, Sarmady M, Procaccio V, Wallace DC. 2013. mtDNA Variation and Analysis Using MITOMAP and MITOMASTER. Curr Protoc Bioinformatics 1: 1.23.1–1.23.26.

Luo S, Valencia CA, Zhang J, Lee N-C, Slone J, Gui B, Wang X, Li Z, Dell S, Brown J, et al. 2018. Biparental inheritance of mitochondrial DNA in humans. Proc Natl Acad Sci USA 115: 13039–13044.

Lutz-Bonengel S, Niederstätter H, Naue J, Koziel R, Yang F, Sänger T, Huber G, Berger C, Pflugradt R, Strobl C, et al. 2021. Evidence for multi-copy Mega-NUMTs in the human genome. Nucleic Acids Res 49: 1517–1531.

Lutz-Bonengel S, Parson W. 2019. No further evidence for paternal leakage of mitochondrial DNA in humans yet. Proc Natl Acad Sci USA 116: 1821–1822.

McCormick EM, Lott MT, Dulik MC, Shen L, Attimonelli M, Vitale O, Karaa A, Bai R, Pineda-Alvarez DE, Singh LN, et al. 2020. Specifications of the ACMG/AMP standards and guidelines for mitochondrial DNA variant interpretation. Hum Mutat 41: 2028–2057.

McKenna A, Hanna M, Banks E, Sivachenko A, Cibulskis K, Kernytsky A, Garimella K, Altshuler D, Gabriel S, Daly M, et al. 2010. The Genome Analysis Toolkit: a MapReduce framework for analyzing next-generation DNA sequencing data. Genome Res 20: 1297–1303.

McLaren W, Gil L, Hunt SE, Riat HS, Ritchie GRS, Thormann A, Flicek P, Cunningham F. 2016. The ensembl variant effect predictor. Genome Biol 17: 122.

Mulero JJ, Fox TD. 1994. Reduced but accurate translation from a mutant AUA initiation codon in the mitochondrial COX2 mRNA of Saccharomyces cerevisiae. Mol Gen Genet 242: 383–390.

Niroula A, Vihinen M. 2016. PON-mt-tRNA: a multifactorial probability-based method for classification of mitochondrial tRNA variations. Nucleic Acids Res 44: 2020–2027.

Perales-Clemente E, Cook AN, Evans JM, Roellinger S, Secreto F, Emmanuele V, Oglesbee D, Mootha VK, Hirano M, Schon EA, et al. 2016. Natural underlying mtDNA heteroplasmy as a potential source of intra-person hiPSC variability. EMBO J 35: 1979–1990.

Preste R, Vitale O, Clima R, Gasparre G, Attimonelli M. 2019. HmtVar: a new resource for human mitochondrial variations and pathogenicity data. Nucleic Acids Res 47: D1202–D1210.

Puttick C, Kumar KR, Davis RL, Pinese M, Thomas DM, Dinger ME, Sue CM, Cowley MJ. 2019. mity: A highly sensitive mitochondrial variant analysis pipeline for whole genome sequencing data. BioRxiv doi: 10.1101/852210.

Romero A, García P. 1991. Initiation of translation at AUC, AUA and AUU codons in Escherichia coli. FEMS Microbiol Lett 68: 325–330.

Shen L, Attimonelli M, Bai R, Lott MT, Wallace DC, Falk MJ, Gai X. 2018. MSeqDR mvTool: A mitochondrial DNA Web and API resource for comprehensive variant annotation, universal nomenclature collation, and reference genome conversion. Hum Mutat 39: 806–810.

Sonney S, Leipzig J, Lott MT, Zhang S, Procaccio V, Wallace DC, Sondheimer N. 2017. Predicting the pathogenicity of novel variants in mitochondrial tRNA with MitoTIP. PLoS Comput Biol 13: e1005867.

Stendel C, Neuhofer C, Floride E, Yuqing S, Ganetzky RD, Park J, Freisinger P, Kornblum C, Kleinle S, Schöls L, et al. 2020. Delineating MT-ATP6-associated disease: From isolated neuropathy to early onset neurodegeneration. Neurol Genet 6: e393.

Stewart JB, Freyer C, Elson JL, Wredenberg A, Cansu Z, Trifunovic A, Larsson N-G. 2008. Strong purifying selection in transmission of mammalian mitochondrial DNA. PLoS Biol 6: e10.

Van Haute L, Powell CA, Minczuk M. 2017. Dealing with an Unconventional Genetic Code in Mitochondria: The Biogenesis and Pathogenic Defects of the 5-Formylcytosine Modification in Mitochondrial tRNAMet. Biomolecules 7: 24.

van Oven M, Kayser M. 2009. Updated comprehensive phylogenetic tree of global human mitochondrial DNA variation. Hum Mutat 30: E386–94.

Wei W, Tuna S, Keogh MJ, Smith KR, Aitman TJ, Beales PL, Bennett DL, Gale DP, Bitner-Glindzicz MAK, Black GC, et al. 2019. Germline selection shapes human mitochondrial DNA diversity. Science 364: eaau6520.

Weissensteiner H, Forer L, Fendt L, Kheirkhah A, Salas A, Kronenberg F, Schoenherr S. 2021. Contamination detection in sequencing studies using the mitochondrial phylogeny. Genome Res 31: 309–316.

Weissensteiner H, Forer L, Fuchsberger C, Schöpf B, Kloss-Brandstätter A, Specht G, Kronenberg F, Schönherr S. 2016a. mtDNA-Server: next-generation sequencing data analysis of human mitochondrial DNA in the cloud. Nucleic Acids Res 44: W64–9.

Weissensteiner H, Pacher D, Kloss-Brandstätter A, Forer L, Specht G, Bandelt H-J, Kronenberg F, Salas A, Schönherr S. 2016b. HaploGrep 2: mitochondrial haplogroup classification in the era of high-throughput sequencing. Nucleic Acids Res 44: W58–63.

Zhang F, Flickinger M, Taliun SAG, InPSYght Psychiatric Genetics Consortium, Abecasis GR, Scott LJ, McCaroll SA, Pato CN, Boehnke M, Kang HM. 2020. Ancestry-agnostic estimation of DNA sample contamination from sequence reads. Genome Res 30: 185–194.

Zhou H, Lin Z, Voges K, Ju C, Gao Z, Bosman LWJ, Ruigrok TJH, Hoebeek FE, De Zeeuw CI, Schonewille M. 2014. Cerebellar modules operate at different frequencies. elife 3: e02536.

